# Synchronous inhibitory pathways create both efficiency and diversity in the retina

**DOI:** 10.1101/214569

**Authors:** Mihai Manu, Lane T. McIntosh, David B. Kastner, Benjamin N. Naecker, Stephen A. Baccus

## Abstract

Visual information is conveyed from the retina to the brain by a diverse set of retinal ganglion cells. Although they have differing nonlinear properties, nearly all ganglion cell receptive fields on average compute a difference in intensity across space and time using a region known as the classical or linear surround^1,2^, a property that improves information transmission about natural visual scenes^3,4^. The spatiotemporal visual features that create this fundamental property have not been quantitatively assigned to specific interneurons. Here we describe a generalizable causal approach using simultaneous intracellular and multielectrode recording to directly measure and manipulate the sensory feature conveyed by a neural pathway to a downstream neuron. Analyzing two inhibitory cell classes, horizontal cells and linear amacrine cells, we find that rather than transmitting different temporal features, the two inhibitory pathways act synchronously to create the salamander ganglion cell surround at different spatial scales. Using these measured visual features and theories of efficient coding, we computed a fitness landscape representing the information transmitted using different weightings of the two inhibitory pathways. This theoretical landscape revealed a ridge that maintains near-optimal information transmission while allowing for receptive field diversity. The ganglion cell population showed a striking match to this prediction, concentrating along this ridge across a wide range of positions using different weightings of amacrine or horizontal cell visual features. These results show how parallel neural pathways synthesize a sensory computation, and why this architecture achieves the potentially competing objectives of high information transmission of individual ganglion cells, and diversity among receptive fields.

---

The parallel architecture of the nervous system makes it challenging to understand the circuit origin of neural computations and the evolutionary advantage of those circuits. Despite new recording and neurostimulation techniques^5,6^, even widely studied computations such as the sensory receptive fields of retinal ganglion cells and orientation selective cells in the visual cortex have not been quantitatively assigned to their neural components^7–9^. Both horizontal and amacrine cells are thought to contribute to the ganglion cell receptive field surround, as indicated by current injection into horizontal cells^10,11^, and pharmacological experiments on amacrine cells, although these latter studies have yielded conflicting results^12–14^. But these results neither define the spatiotemporal feature contributed by a particular interneuron to the ganglion cell linear receptive field, nor reveal the functional benefits of utilizing such a parallel architecture.

We first sought to directly measure the spatiotemporal contributions of individual interneurons to the linear ganglion cell receptive field surround. The sensory feature, *C*_*x,t*_, contributed by an interneuron to a downstream neuron is created in two stages—first the transformation from the stimulus to the interneuron, which is the interneuron’s own spatiotemporal receptive field, *F*_*x,t*_—and second the transformation, *G*_t_, between the interneuron and the downstream neuron in its projective field^15–17^ (Fig. 1A). The combined effect of these two functions has not been measured, and thus the contributions of individual interneurons are unknown. To measure the receptive field component, *C*^*(a)*^, contributed by single sustained amacrine cells, which have linear responses, ^16,18^ to the ganglion cell receptive field, we presented a one-dimensional spatiotemporal stimulus consisting of randomly flickering lines. We first focused on the time course of *C*^*(a)*^ by computing the spatial average, 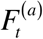, of the amacrine cell’s receptive field (Fig. 1–BD). Next, we computed the temporal filter, 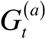, describing the transmission from amacrine to ganglion cell by injecting white noise current for 300 s into the amacrine cell and correlating that current with the recorded ganglion cell spikes. This transmission filter 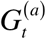 had a monophasic negative peak (Fig. 1E), indicating that the amacrine cell was inhibitory^18^.

**Figure 1.**
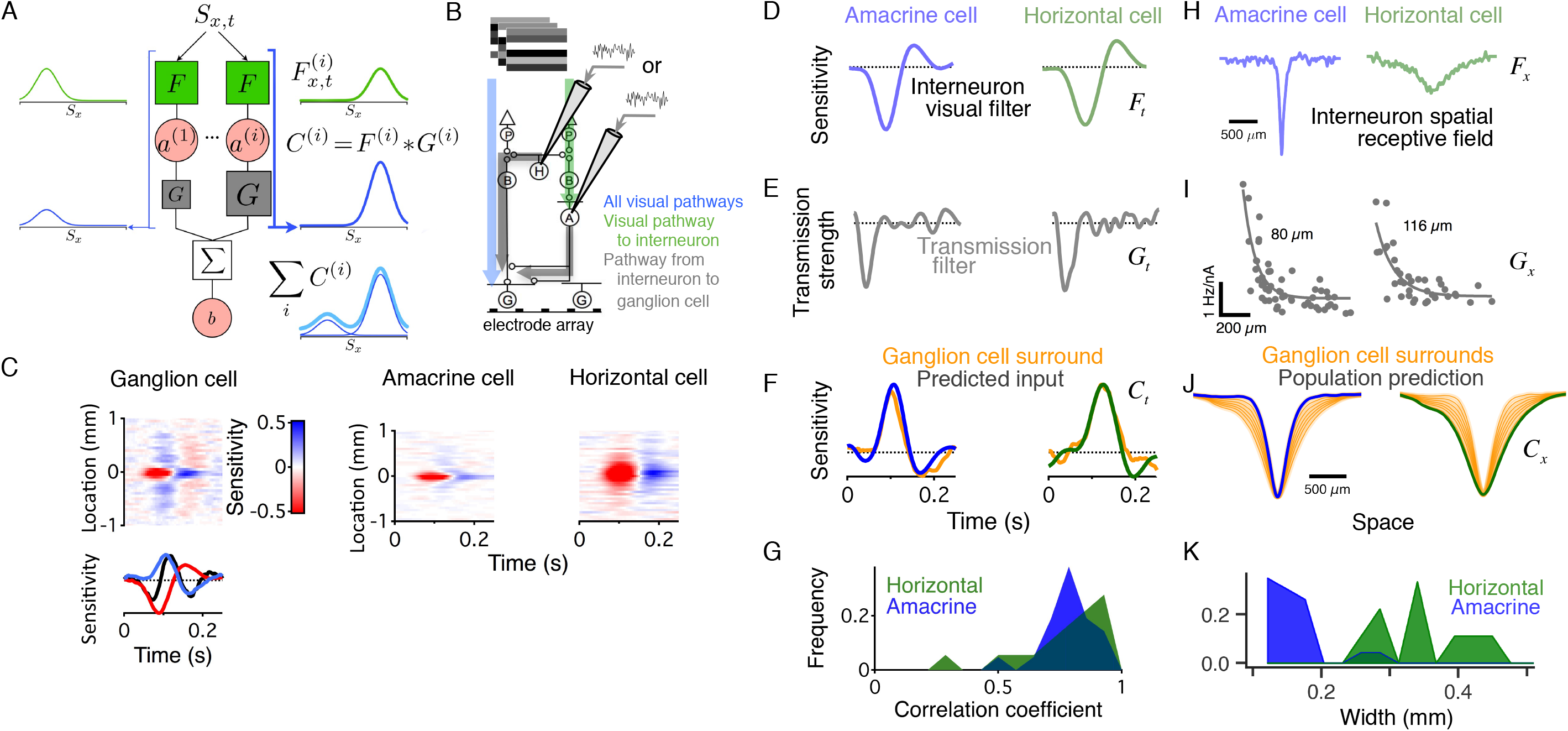
Synchronous visual features from inhibitory pathways at different spatial scales. A) Schematic of a hypothetical linear neuron that receives input from multiple neural pathways. The response of each interneuron 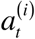 is the convolution of the stimulus and its particular linear spatiotemporal filter 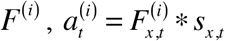, where * indicates a convolution. The response of the interneuron *b* is the sum of the outputs of 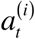, each filtered through a transmission filter 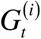, such that 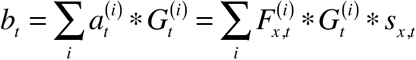. The visual feature *C*^*(i)*^ contributed each interneuron *a*_*i*_ is a combination of the neural pathways leading into and out of *a*_*i*_ (blue bold lines for one interneuron), i.e. a convolution of the spatiotemporal filter, *F*^*(i)*^, from the stimulus to interneuron *a*_*i*_, and the temporal filter, *G*^(*i*)^ from *a_i_* to *b*. B) Experimental arrangement showing simultaneous intracellular and multielectrode recording. Colored arrows correspond to measured temporal filters shown in F–G. C) Top left. One-dimensional spatiotemporal map of a fast Off-type retinal ganglion cell. Bottom. Spatial average of receptive field surround (blue), center (red) and total receptive field (black). Right. One-dimensional receptive field of an amacrine cell and a horizontal cell. D) Spatial average of amacrine cell, 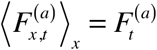 and horizontal cell. E) Left. Transmission filter, 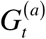, between amacrine and ganglion cell computed from white noise current injection in the presence of the flickering lines visual stimulus to place the ganglion cell in a similar state of adaptation as in the control condition. Right. The transmission filter 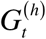 for a horizontal cell. F) Left. The visual feature, 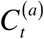 conveyed by the amacrine cell, computed as the convolution between 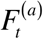 and 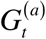, corrected for the membrane time constant, compared with that of the ganglion cell surround computed by spatially averaging the region that lay outside the receptive field center. Right. The same for a horizontal cell and different ganglion cell. G) Histogram of correlation coefficients between predicted interneuron transmission and ganglion cell surround time course. H) Spatial one-dimensional receptive field of amacrine and horizontal cells. I). The strength of transmission between amacrine cells (left) or horizontal cells (right) and ganglion cells computed from the average slope of a nonlinearity computed during white noise current injection into the interneuron. The nonlinearity was taken from a linear-nonlinear (LN) model computed between each cell pair during current injection^18^. J) Amacrine and horizontal cell population predictions estimated by convolving the interneuron receptive fields with their appropriate spatial transmission filters, compared to retinal ganglion cell receptive field spatial surrounds estimated from the linear receptive field model in Fig 3C. K) Histogram of the sizes of a population of linear amacrine and horizontal cells, measured as one standard deviation of a Gaussian fit.

We then estimated the temporal feature conveyed by the amacrine cell to the ganglion cell as 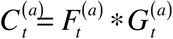, convolving the amacrine cell temporal receptive field, 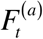, with the amacrine to ganglion cell transmission filter, 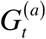, correcting for a double contribution of the amacrine cell membrane time constant^18^ (see methods). The visual feature 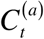 conveyed by this amacrine cell to the ganglion cell was an increase in light intensity with a latency of ~120 ms.

We found that 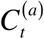 matched very closely the time course of the ganglion cell spatial surround (Fig. 1F). The correlation coefficient, *r*, was 0.81 ± 0.02 (Fig.1G, n = 21 cell pairs), similar to the measured variation within the ganglion cell surround itself, computed between the two opposite sides of the ganglion cell surround (r = 0.83 ± 0.03).

The same analysis was then performed with horizontal cells rather than amacrine cells (Fig. 1D–G). Although one might expect that different interneurons with different temporal kernels would convey different temporal features, we instead found that the visual feature contributed by a horizontal cell to ganglion cells, 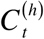, also matched the time course of the ganglion cell surround for each cell pair (*r* = 0.81 ± 0.04, n = 18 cell pairs). Thus, the visual features conveyed through the distinct neural pathways of amacrine and horizontal cells were synchronous in time, and closely matched the time course of the ganglion cell receptive field surround.

We then estimated the spatial receptive field components 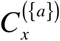 and 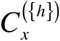 contributed by amacrine and horizontal cell populations to a single ganglion cell. Because each cell type tiles the retina in space^19^, the spatial weighting of one amacrine cell’s projective field^16^—equivalent to the amacrine to ganglion cell point spread function—is equivalent to the spatial weighting 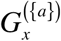 of the convergence of many amacrine cells to one ganglion cell. We thus convolved the spatial receptive field (Fig. 1H) of each interneuron class with its measured projective field (Fig. 1I) to estimate the spatial visual feature conveyed by a population of amacrine or horizontal cells to a single ganglion cell (Fig. 1J). The receptive field component conveyed by the amacrine cell population on average had a smaller half-maximal width (332 μm +/− 75 μm s.d.) than that of horizontal cells (740 μm +/− 139 μm s.d.), with the surround sizes of ganglion cells falling in a range in between (Fig. 1 J, K). Thus amacrine and horizontal cells conveyed the same temporal feature at different spatial scales that spanned the range of sizes of ganglion cell receptive field surrounds.

The above analysis relied on a linear model of the transformations from stimulus to interneuron to ganglion cell, and furthermore assumes that the interneuron’s effects were the same under perturbation by white noise current injection as during visual input. To measure the interneuron’s contribution without these assumptions, we designed a direct causal test to measure whether the interneuron’s timed visual responses specifically generated the ganglion cell surround. We used a full field visual stimulus to measure the temporal receptive field of the ganglion cell. Although this measurement sums over the spatial center and surround of the cell, the first peak in the ganglion cell temporal receptive field receives little contribution from the spatial surround, and the second opposing peak is almost exclusively comprised by the spatial surround (Fig. 1C, Extended Data Fig. 1).

To directly test whether amacrine transmission contributed to the ganglion cell temporal surround, we amplified or diminished the amacrine cell’s visually driven voltage fluctuations (Fig. 2A). We first recorded ganglion cell and amacrine cell responses to visual stimuli alone without current, then played back timed current that either amplified or diminished the voltage fluctuations of the interneuron while repeating the visual stimulus. This record and playback method perturbed the cell only at the times of the visually driven response, avoiding potential off-target effects created by mistimed perturbations^18^.

**Figure 2.**
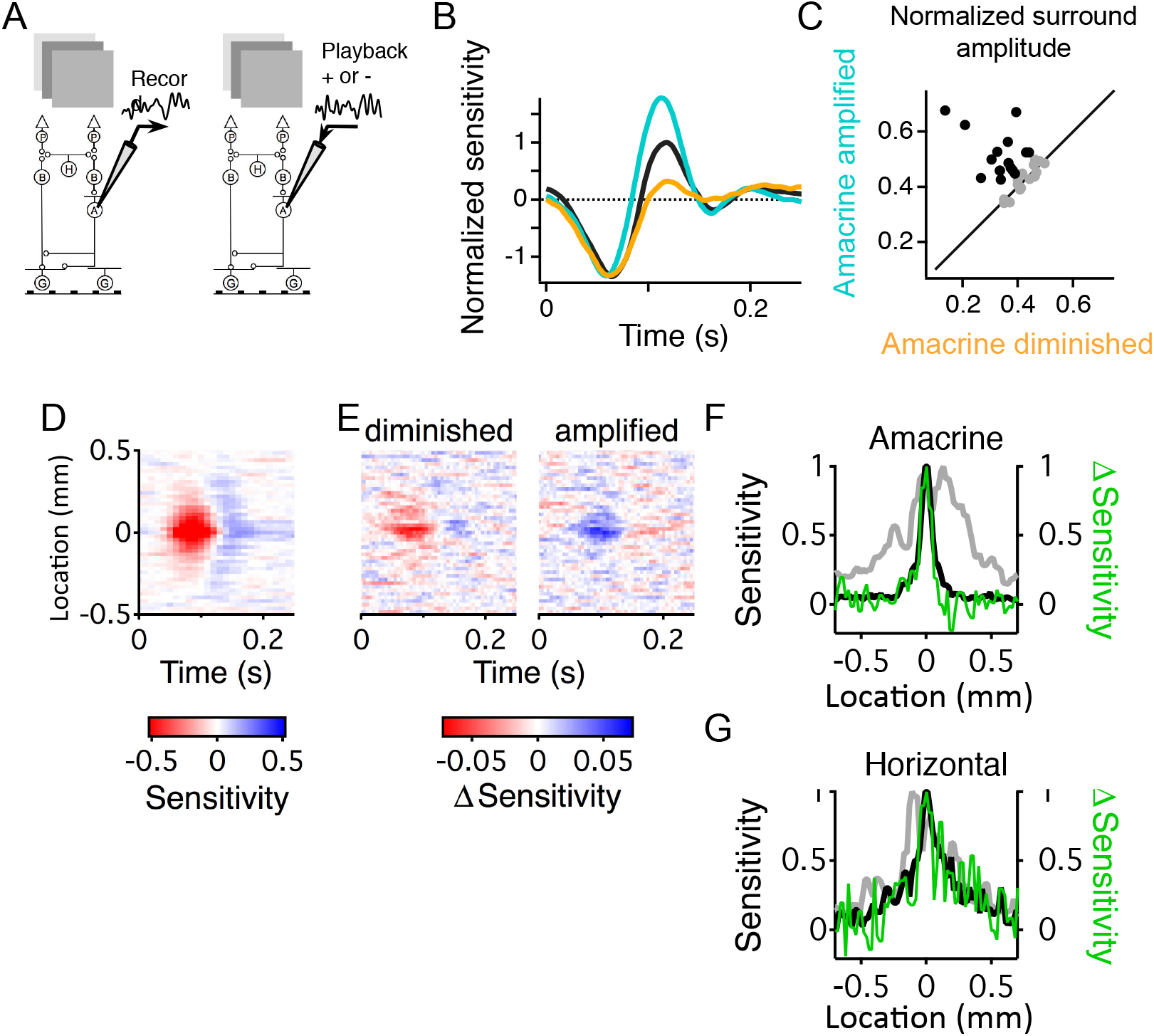
Direct causal measurements of the visual feature conveyed by an interneuron. A) Schematic of experiment recording the membrane potential response of an amacrine cell. The visual stimulus was then repeated along with current injection timed to either amplify or diminish the amacrine cell’s visually driven membrane potential fluctuations (see methods). B) Visual sensitivity to a uniform visual stimulus computed in the control condition, with amacrine transmission amplified, and with amacrine transmission diminished. Filters were scaled in amplitude so that all sensitivity was represented in the filter (see methods). C) Amplitude of the surround compared for conditions when amacrine transmission was amplified or diminished (n = 6 amacrine cells and 31 ganglion cells). Black symbols indicate amacrine-ganglion cell pairs for which a monophasic or biphasic transmission filter could be computed from the separate protocol of white noise current injection (Fig. 1). D) One-dimensional spatiotemporal receptive field of an amacrine cell. E) Change in linear receptive field averaged across five ganglion cells when the amacrine cell’s output was amplified (right) or diminished (left) using record and playback. Ganglion cells were chosen whose receptive fields overlapped that of the amacrine cell. Results shown for a single amacrine cell (n = 5 ganglion cells) F) Spatial amplitude of the amacrine cell receptive field (black) compared with the difference in visual sensitivity across a population of ganglion cells between when amacrine cell fluctuations were amplified or diminished (green) (10 amacrine cells and 70 ganglion cell pairs). A Gaussian fit to the average amacrine cell receptive field had a standard deviation of 56 μm, smaller than the total region of visual sensitivity resulting from summed ganglion cell receptive fields, which was 336 μm. Spatial extent of recorded ganglion cells shown in grey. All curves normalized by their maximum value for ease of spatial comparison. G) Same as F) for horizontal cells (3 horizontal cells and 18 ganglion cells). A Gaussian fit to the average horizontal cell receptive field had a standard deviation of 283 μm, smaller than the total region of visual sensitivity of the measured ganglion cell population, which was 363 μm.

Amplifying the amacrine cell’s output increased the amplitude of the ganglion cell temporal surround, with only a very small change in the sensitivity of the negative peak, which derived from the receptive field center (Fig. 2 B, C). Conversely, canceling the amacrine cell’s visually driven voltage fluctuations in some cases caused the ganglion cell’s temporal surround to nearly disappear, whereas the effect on the first peak was minor. These results show causally that the visual feature conveyed by amacrine cells is used to construct the ganglion cell temporal surround.

The above results imply that a linear amacrine cell contributes a visual feature equal to its own linear receptive field filtered through a temporal synaptic delay. We tested this idea directly by amplifying or diminishing the voltage fluctuations of an amacrine cell during a one-dimensional spatiotemporal stimulus. Comparing the amacrine amplified and diminished conditions showed that the ganglion cell population experienced a localized reduction in the receptive field that spatially matched the amacrine cell’s receptive field (Fig. 2D–F). This demonstrates causally that amacrine cells contribute their own receptive field to construct the ganglion cell receptive field. Note that this result is not guaranteed, and that nonlinearities such as a multiplicative interaction between neural pathways could cause an interneuron to deliver a visual feature that is the combination of its own receptive field and other pathways (Extended Data Fig. 3, Supplementary Information).

Similar experiments using the record and playback technique applied to horizontal cells altered the spatiotemporal receptive field of ganglion cells. As with amacrine cells, the change in sensitivity across the ganglion cell population matched the horizontal cell spatial receptive field (Fig. 2G). Thus, the visual feature comprising the ganglion cell surround was not an average of differently timed contributions from separate interneuron pathways. Instead, two distinct pathways contributed the same temporal feature that matched the final ganglion cell surround, yet at different spatial scales.

What is the advantage of having two neural pathways conveying the same temporal visual feature at different spatial scales to the ganglion cell linear receptive field? Ganglion cell receptive fields approximately maximize information transmission given the statistics of natural visual images ^3,4,20,21^. We confirmed that the average ganglion cell receptive field was consistent with these previous claims (Fig. 3A, Extended Data Fig. 2). Yet ganglion cells have different receptive field shapes ^19,22,23^ (Fig. 3B) to support different functional roles such asymmetric receptive fields in direction selectivity^24^ or selecting for particular speeds of motion^25^. Furthermore, it has been proposed that diverse neural responses reduce neural correlations and increase the information transmission capacity of a neural population^26–34^. If there is one optimal shape of a receptive field, one would expect that a diverse population must have some cells with suboptimal information transmission. This tradeoff between diversity and optimal information transmission is not well understood.

We modeled the ganglion cell receptive field as a linear combination of the measured average visual features contributed by horizontal and amacrine cells, as well as an excitatory central region (Fig. 3C). This model captured 93% ± 0.3% (n = 1382) of the variance of measured ganglion cell receptive fields, indicating that we have accounted for the main interneuron contributions that create the linear surround (Extended Data Fig. 4). Based on measured noise in fast Off ganglion cells, and photoreceptor noise estimates based on the previously measured mean vesicle release rate (see methods) we computed a fitness landscape of information transmission for each type of receptive field. This analysis revealed that around the single optimal receptive field was an extended ridge of receptive field shapes with near-optimal information transmission. This near-optimal region was achieved when the horizontal and amacrine cell weighting caused the center weight to slightly exceed that of the surround (Fig. 3D, E). Thus, the measured components of horizontal and amacrine cells together define a direction within this landscape in which receptive fields can be changed without impacting information transmission.

**Figure 3.**
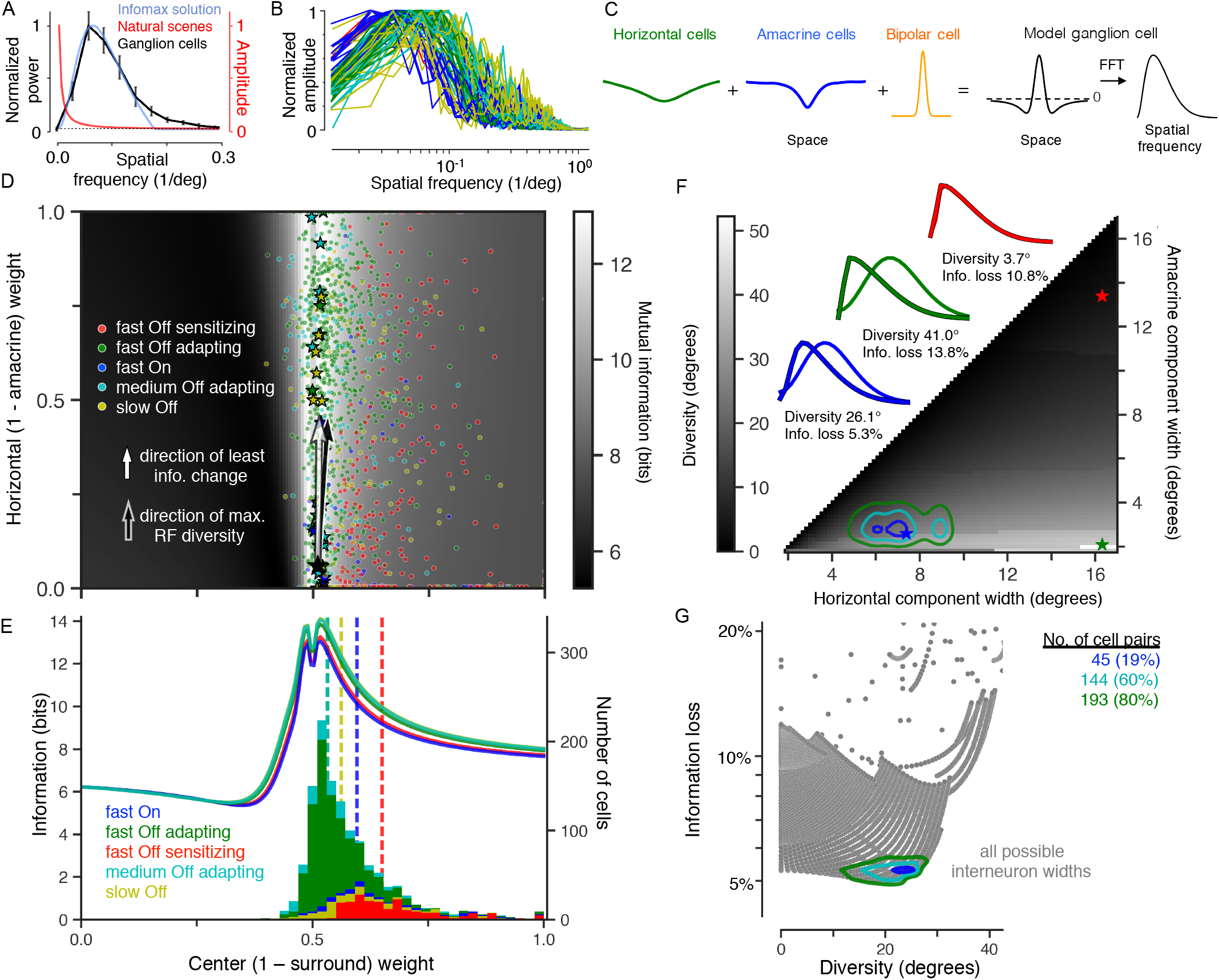
Horizontal and amacrine cell contributions create diverse retinal ganglion cell receptive fields that maximize information transmission. A) Normalized power spectra of the average retinal ganglion cell receptive field (n = 13, black) and the linear filter that maximizes information about natural scenes (blue), together with the amplitude spectrum of natural scenes (red). B) Example retinal ganglion cell receptive fields in the frequency domain, colored by cell type. C) Ganglion cell linear receptive field modeled as a linear combination of horizontal and amacrine cell populations, together with a Gaussian center. D) Greyscale map shows the information fitness landscape, computed as the mutual information between natural scenes and the output of linear receptive field models as a function of different horizontal and amacrine weightings (ordinate) and different excitatory center contributions (abscissa). Colored symbols indicate 1,354 ganglion cell receptive fields of different cell types, 772 fast Off adapting, 217 fast Off sensitizing, 159 medium Off, 58 fast On and 148 slow Off, parameterized by the model in (C), and stars are the examples in (B). Large black star at the origin of the two vectors indicates the single ideal receptive field, which lies on the ridge of near-optimal information transmission. Light arrow indicates the direction of least information loss with respect to the ideal receptive field. Dark arrow indicates the direction of largest variation of ganglion cell receptive fields, computed using principal components analysis. E) Information profile for different cell types as a function of receptive field center weight. Curves show information transmission at an interneuron weighting of 0.5 horizontal cell weight, which is a horizontal slice through the grayscale image in (D). Stacked histograms show the number of cells with receptive fields at each center weighting. Dotted lines denote the medians of the distributions. The slight trough at equal center-surround weighting (0.5) reflects the loss of information when the mean intensity is completely rejected. F) Heat map of the angle between a receptive field with no amacrine component or no horizontal component, as a function of the width of both components. Along the diagonal the pathways are identical and create zero diversity. The maximum achievable diversity drops off as both receptive fields become large. Three examples of the maximally diverse receptive fields are plotted in the spatial frequency domain, with the parameters for these examples denoted by stars. Contours denote where pairs of recorded horizontal and amacrine cells lie on this plot, computed for 9 horizontal cells and 27 amacrine cells whose receptive field population component ranged in one standard deviation width from 256 to 453 μm (horizontal cells) and 108 to 284 μm (amacrine cells) in diameter. G) The fractional information loss relative to the ideal receptive field plotted against the maximal ganglion cell diversity achievable with different weightings of horizontal and amacrine cells for all possible models from (D) with different interneuron component widths. Contours indicate where the population of horizontal and amacrine cells lie on this plot.

When we examined the actual receptive fields of over 1,300 ganglion cells, their receptive fields closely approximated this ridge of near-optimal transmission yet had a broad distribution of receptive fields types, ranging from having surrounds matching the horizontal cell feature with no amacrine contribution to those with only an amacrine contribution. Individual cell types had different median values of amacrine contribution, yet varied broadly within cell type (Extended Data Fig. 5). Although some cell types had weaker surrounds than the optimal value, cell types with higher noise systematically had weaker surrounds consistent with theories of information maximization^3,4^ (Extended Data Fig. 6). Within this landscape, we computed the direction of greatest variation of receptive field shape across the ganglion cell population, and found it was nearly identical to the direction of least loss of information transmission, differing by ~2 degrees (Fig. 3D). This striking correspondence between the fitness landscape that maximizes information and the diversity of ganglion cell receptive fields indicates that evolution has generated receptive field diversity in a direction of neutral impact to information efficiency. Although it is unknown how this remarkable relative tuning of amacrine and horizontal cell input is achieved, because the horizontal cell surround component is already subtracted from the amacrine cell receptive field, a plausible explanation is that adding a greater weighting of amacrine cells automatically reduces the horizontal cell contribution.

To consider the effects of different amacrine cell types on a more general ganglion cell population, we then analyzed how a measured inhibitory interneuron population (n = 36) would influence the two separate properties of information transmission and diversity. Taking all possible amacrine and horizontal cell pairs, we found that the inhibitory interneuron population is positioned at a point that maximizes receptive field diversity in the ganglion cell population while minimizing information loss due to a suboptimal receptive field (Fig. 3F–G). Interneuron pathways with the spatial scales of amacrine and horizontal cells support the construction of a diverse population of ganglion cell surrounds, each of which is near-optimal in terms of information transmission under natural scene statistics.

These results offer a quantitative and theoretical explanation as to how and why two parallel inhibitory pathways generate the ganglion cell classical receptive field. Our analysis quantitatively accounts for the measured optimal linear receptive fields of ganglion cells, and explains the structure of their observed diversity. Although the question of what receptive field properties are optimal has been studied^3,4,20,35–37^, little attention has been given to how neural circuits should generate those receptive fields. Linear computations pose an added difficulty, as neural pathways that carry the same signals and are summed without distortion cannot be separately identified without a selective way to perturb each component. In contrast, parallel pathways with distinct nonlinearities have enabled recent theoretical and experimental studies to reveal the benefits of separate neural pathways differing in their thresholds^31,32^.

Our approach to identify the contribution of a cell to a neural function can be applied to more complex nonlinear computations, other stimulus modalities and optogenetic perturbations. Critical to this process will be to measure the composition of input and output functions by both recording from a neuron and perturbing it in order to avoid misinterpretations that arise from optogenetic perturbation alone^38^. In the case of retinal receptive fields, this approach reveals how multiple interneuron pathways in the retina’s parallel and layered circuitry maintain an efficient representation, while allowing the evolution of a diverse neural population.

## Materials and Methods

### Visual stimuli

Stimuli were projected from a video monitor at a photopic mean intensity of 10 mW/m^2^, and were drawn from a Gaussian distribution unless otherwise noted. The contrast of stimuli defined as standard deviation of the intensity distribution divided by the mean ranged from 10 - 35 %.

### Simultaneous intracellular and multielectrode recording

Methods for simultaneous intracellular and extracellular recording using a 60-electrode array in the intact salamander retina were as described^18^. Briefly, intracellular electrodes (150 - 250 Μ**Ω**) were used for either recording or current injection in bridge mode. To compute the temporal receptive field component contributed by an amacrine or horizontal cell, we computed a visual filter *F*_*t*_ as the spatial average of the spatiotemporal receptive field *F*_*x,t*_ between the visual stimulus and the interneuron membrane potential. We then computed a transmission filter *G*_*t*_ between white noise current injected into the cell and ganglion cell spikes. Current amplitudes were chosen so that they were estimated to maintain the membrane potential within a physiological range (~10 mV s. d.) given an estimate of the membrane conductance measured using pulses of current^18^. This value was 0.5 nA s.d. for amacrine cells and 0.5 − 1.0 nA s.d. for horizontal cells. To compute the predicted transmission, *C*_*t*_ for a neural pathway as a composition of *F*_*t*_ and *G*_*t*_, because *F*_*t*_ was computed between the stimulus and interneuron membrane potential, but *G*_*t*_ was computed from injected current, we corrected for the double contribution of the amacrine cell membrane time constant *τ* by deconvolving with the function *e*^−*t*/*τ*^ as previously described^18^.

For record and playback experiments, to amplify or diminish the fluctuations of an interneuron, first the membrane potential fluctuations were recorded without current and an exponential function representing the membrane time constant of the cell was measured. The recorded membrane potential fluctuations were then deconvolved by the exponential function, creating a current sequence that when filtered according to the membrane time constant was predicted to match the measured voltage response. With cells that have slower membrane potential fluctuations such as sustained amacrine and horizontal cells, this procedure is not highly sensitive to an accurate measurement of the membrane time constant^18^. To amplify the cell’s voltage fluctuations, the visual stimulus *s(t*) was repeated while injecting the current *I*_*a*_ (*t*) synchronized with the visual stimulus so as to make both depolarizations and hyperpolarizations larger. The standard deviation of the current was set to 500 pA for amacrine cells and 750 pA for horizontal cells. To diminish the cell’s voltage fluctuations, *s*(*t*) was repeated while injecting the current − *I*_*a*_ (*t*), thus partially canceling the cell’s visual input. This allowed us to compare two opposite perturbations of the input.

Sustained Off-type amacrine cells likely comprise multiple cell types, most of which have a narrow receptive field (< 200 μm), and were identified by their sustained, linear flash responses, the presence of an inhibitory surround, and their inhibitory transmission to Off-type ganglion cells^18^. Horizontal cells were identified by their lack of receptive field surround, linear response, and a receptive field center that exceeded 200 μm in diameter. Ganglion cells were classified by a white noise stimulus as described^39^. Fast Off-type ganglion cells include two distinct cell types, adapting and sensitizing that form independent mosaics, but here they were analyzed together unless otherwise noted.

### Linear model of visual responses and interneuron transmission

Linear models of visual responses, and of amacrine and horizontal cell transmission were computed as described using the standard method of reverse correlation^18^. For current injection, the stimulus, *i* (τ), was white noise current (bandwidth of 0 - 50 Hz), and was convolved with a linear temporal filter, *F*_*t*_ = *F*(*t*), which was computed as the time reverse of the spike triggered average stimulus, such that

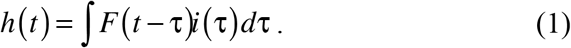

To compare the absolute sensitivity in spatial and temporal regions of the receptive field between conditions, all sensitivity was placed in the linear filter. To do this, the linear filter was extended to a linear-nonlinear (LN) model by computing a static nonlinearity *N(h)* that captured the threshold and average sensitivity of the cell. The nonlinearity was then scaled along the input axis in the condition of current injection so that *N(h)* was the same as in the control condition, and the linear filter was scaled by the same factor on the vertical axis^21^. This procedure left the overall LN model the same, but placed all sensitivity in the linear filter.

Spatiotemporal receptive fields were measured using reverse correlation^40^ of the firing rate or membrane potential response with a visual stimulus consisting of independently modulated 100 μm squares or 50 μm wide bars.

The normalized contribution of the surround (Extended Data Fig. 1) was computed as

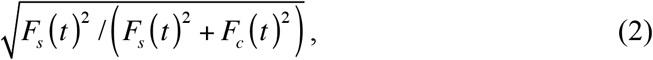

where *F*_*s*_ is the time course of the linear receptive field averaged over the spatial surround, and *F*_*c*_ is the time course of the center. This measure varies between zero (no surround contribution) and one. Results were combined across cells by stretching the total temporal receptive field *F*_*t*_ in time to align the negative and positive peaks. The center time course was computed as *F*_*c*_ (*t*) = *F*_*t*_ (*t*) − *F*_*s*_ (*t*).

### Optimal receptive fields

The ideal spatial linear ganglion cell receptive field was taken to be one that maximizes information transmission about natural scenes. We solved for this optimal receptive field in the frequency domain assuming constrained ganglion cell response variance^3^. This problem can be reformulated as minimizing redundancy, *min*_*F*_ *C*(*Y*) − *λI*(*X;Y*), where *I*(*X;Y*) is the mutual information between the retina’s visual input *X* and ganglion cell response *Y* = *F*(*X* + *N*_*in*_) + *N*_*out*_, where *N*_*in*_ is the input noise in the retinal circuit prior to the ganglion cell,*N*_*out*_ is the output noise that corrupts the ganglion cell response after the stimulus is filtered by the linear receptive field, and *C*(*•*) is the channel capacity, or upper bound on the information the optimized linear receptive field can transmit^3,20,35^. Constraining the receptive field *F* to be linear and spatially symmetric, and making a Gaussian approximation assumption on the visual stimulus *XF*, we can compute the optimal ganglion cell receptive field *F* explicitly. The amplitude spectrum, *F* = *F* (ω) of the spatial receptive field is given by

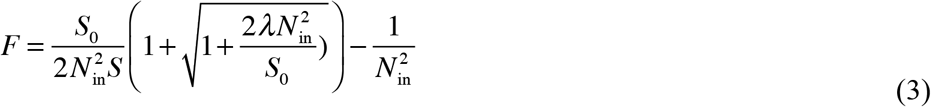

where ω is spatial frequency, S_0_ (ω) is the power spectrum of the visual input *X*, 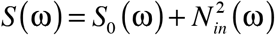 is the power spectrum of the signal + noise input the ganglion cell receives, and the Lagrange multiplier λ is found by numerically integrating the following expression,

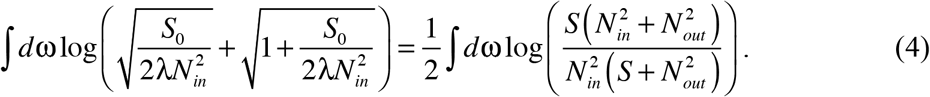

The spatial receptive field *F* that maximizes information transmission depends only on the signal power spectrum and the input and output noise amplitude spectra. For all optimal receptive fields in this paper we used the average power spectrum of natural images obtained from a database^41^, and fixed the input and output noise to be Gaussian white noise. We constrained the total signal-to-noise ratio (SNR), var(*X*)/var(*XN*_*in*_ + *N*_*out*_), to match the average SNR estimated from the trial-to-trial variability of 28 fast Off ganglion cells simultaneously recorded in response to a repeated 35% contrast natural scenes sequence. The SNR in these experiments was measured by dividing the variance of each trial, averaged over all trials, by the variance across trials averaged across time. The relevant cone photoreceptor noise is that which occurs prior to spatial filtering by horizontal cells, and includes photoreceptor vesicle release. To estimate the SNR, we assumed a mean vesicle release rate^42^ of 750 s^−1^, Poisson noise, and a 100 ms integration time, meaning that in one integration time an average of 75 vesicles are released. We then computed the SNR as a function of the temporal contrast of a Gaussian stimulus, defined as the standard deviation divided by the mean intensity (Extended Data Fig. 2). An average value of contrast of 0.3 previously reported for natural scenes^43^, corresponds to an SNR of ~7. We also computed how information transmission changed over a range of SNR values, (Extended Data Fig. 7).

To investigate the contributions of the horizontal cell and linear, or sustained, amacrine cell populations, *C*_{*h*}_ and *C*_{*a*}_ to the ganglion cell linear receptive field, we convolved the average amacrine cell spatial receptive field 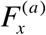 with the amacrine cell projective field 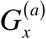 and similarly convolved the horizontal cell receptive and projective fields, and 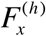 and 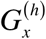 (Fig. 1D), respectively. This convolution represents the spatial weighting of the horizontal and amacrine population contributions to a given ganglion cell, assuming that both interneuron and ganglion cell populations tile the retina.

To explore how well horizontal or amacrine cell populations alone could contribute to a ganglion cell surround that maximizes information, we modeled the ganglion cell spatial receptive field as the linear combination *αB*^*μ,σ*^ +(1 −*α*)(*ηC*_*{h}*_ +(1−*η*)*C*_*{a}*_), where *α* ∈ [0,1] represents the relative weight between center and surround, and *η* ∈ [0,1] represents the relative weight between horizontal and amacrine contributions. For the amacrine-only surround, *η* = 0, and for the horizontal-only surround, *η =* 1. The center contribution *Β*_*σ*_ is a Gaussian where both the mean *μ* and standard deviation σ are jointly fit with *η,#x03B1;*. Two parameters allowed for a spatial offset of the horizontal and amacrine receptive field components to account for the observation that center and surround were not always perfectly concentric. When fitting individual retinal ganglion cells (Fig. 3C–F), parameters were refit for each cell. The information landscape in Fig. 3D did not change qualitatively when *σ* was chosen to be the average center width of the five different retinal ganglion cell types.

To enforce that the weights *η* and *α* were between 0 and 1 and also maintain a smooth gradient for optimization, *η* and *α* were defined as *η* = *θ*(*η*′) and *α* = *θ*(*α*′), where *θ*() is a sigmoidal function with a range between 0 and 1. The alternate parameters *η*′ and *α′* were then optimized. We verified using simulated data that the fitting procedure could recover weights *η* and *α* that were exactly 0 or 1.

### Diversity vs. information loss

In Figure 3F–G, diversity was computed by calculating the angle between model ganglion cells with no amacrine cell contribution or no horizontal cell contribution. The angle between these models is defined as 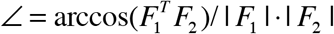. To account for these effects across an interneuron population, these calculations were repeated with each horizontal-amacrine pair using 9 horizontal cells, and 27 different amacrine cells. Information loss was defined as the percentage decrease in information transmission relative to the maximum possible information transmission achievable with any combination of model parameters.

## Supplementary Information

Is the result that an interneuron contributes its own linear receptive field to the linear receptive field of a downstream neuron the only possibility, or are there other potential results that depend on nonlinearities in the circuit? To demonstrate that this result is not guaranteed, it is sufficient to give a theoretical example where changing the gain of an interneuron as in the experiment in Fig. 2 can change the receptive field of a downstream neuron *b* in a manner different from the receptive field of *a*_1_. As a simple example, take a neuron *b* that responds to two independent stimuli *s*_1_ and *s*_2_, each with a Gaussian distribution, zero mean. Responses are instantaneous, so time can be ignored. Two interneurons - with single region receptive field *s*_1_, and *a*_2_ with receptive field *s_2_* - contribute to b’s response. The response of *b* is

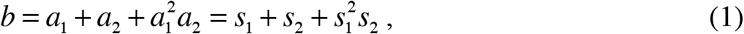

meaning that the output neuron *b* sums over *a*_1_ and *a*_2_, but there also a multiplicative component that combines 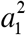 and *a*_2_. The Record and Playback experiment in Figure 3 considered here would change the gain of by a factor *γ*, such that 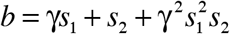. The receptive field *D* of *b* is a two-component vector, one each for *s*_1_ and *s*_2_, and can be computed by the standard method of correlating the stimulus with the response, normalized by the stimulus autocorrelation. The standard equation to compute the receptive field for a white noise input is,

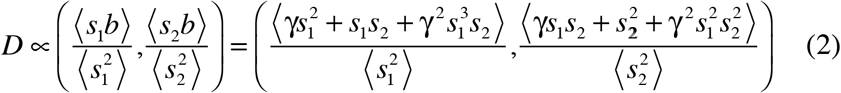

where brackets, 〈…〉, denote the average over stimuli. Because *s*_1_ and *s_2_* have zero mean and are independent, this equation reduces to,

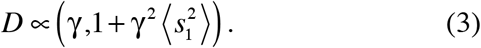

The question of how the receptive field *D* changes when the amplitude of *a*_1_ is changed by the factor *γ* is equivalent to the partial derivative of *D* with respect to *γ*. This is equal to

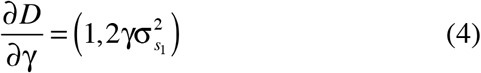

where 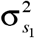 is the variance of *s*_1_. Thus, the gain of *a*_1_ changes the receptive field of b not only by the receptive field of *a*_1_, which is (1,0), but the receptive field component contributed by *a*_1_ also includes the receptive field of *a*_2_. It is the multiplicative term 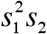 in equation (1) that causes this effect, because the correlation of *s_2_* yields the term 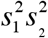 which has non-zero mean. Thus, the result of Fig. 2, where each interneuron contributes its own linear receptive field to the downstream ganglion cell is not guaranteed.

More generally, one can consider a neuron *b* that is described by a polynomial function of interneuron inputs *a*_1_ and *a*_2_, and a stimulus direction *s_2_* that contributes to the receptive field of *a*_2_ but not *a*_1_. If the polynomial expression of *b* contains a term 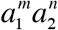 where *m* is even and *n* is odd, then the correlation of a stimulus direction *s*_2_ with *b* will yield a term with an even power of both *a*_1_ and *a*_2_, which will have a non-zero mean. Therefore, changing the gain of the *a*_1_ will also change the contribution of *s*_2_ to *b*, even though *s*_2_ does not contribute to the receptive field of *a*_1_.

**Extended Data Figure 1.**
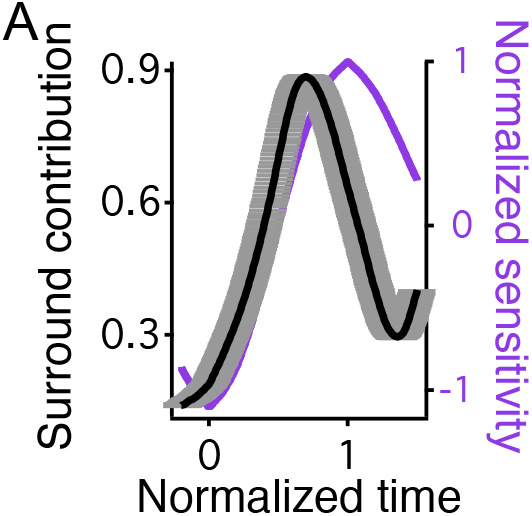
Contribution of spatial surround to the total ganglion cell temporal filter. Ganglion cell spatiotemporal receptive fields were mapped in one dimension. Shown is the normalized ratio (see methods) between the amplitude of the linear receptive field averaged over the spatial surround region, and the amplitude averaged over the entire receptive field. This ratio shows the contribution at each time point of the surround to the total receptive field averaged over space. Thin line indicates the mean, thick line indicates mean ± s.e.m. Results are combined for 1003 fast off-type ganglion cells by stretching each filter in time to align both the negative peak (shown at t = 0) and positive peak (shown at t = 1).

**Extended Data Figure 2.**
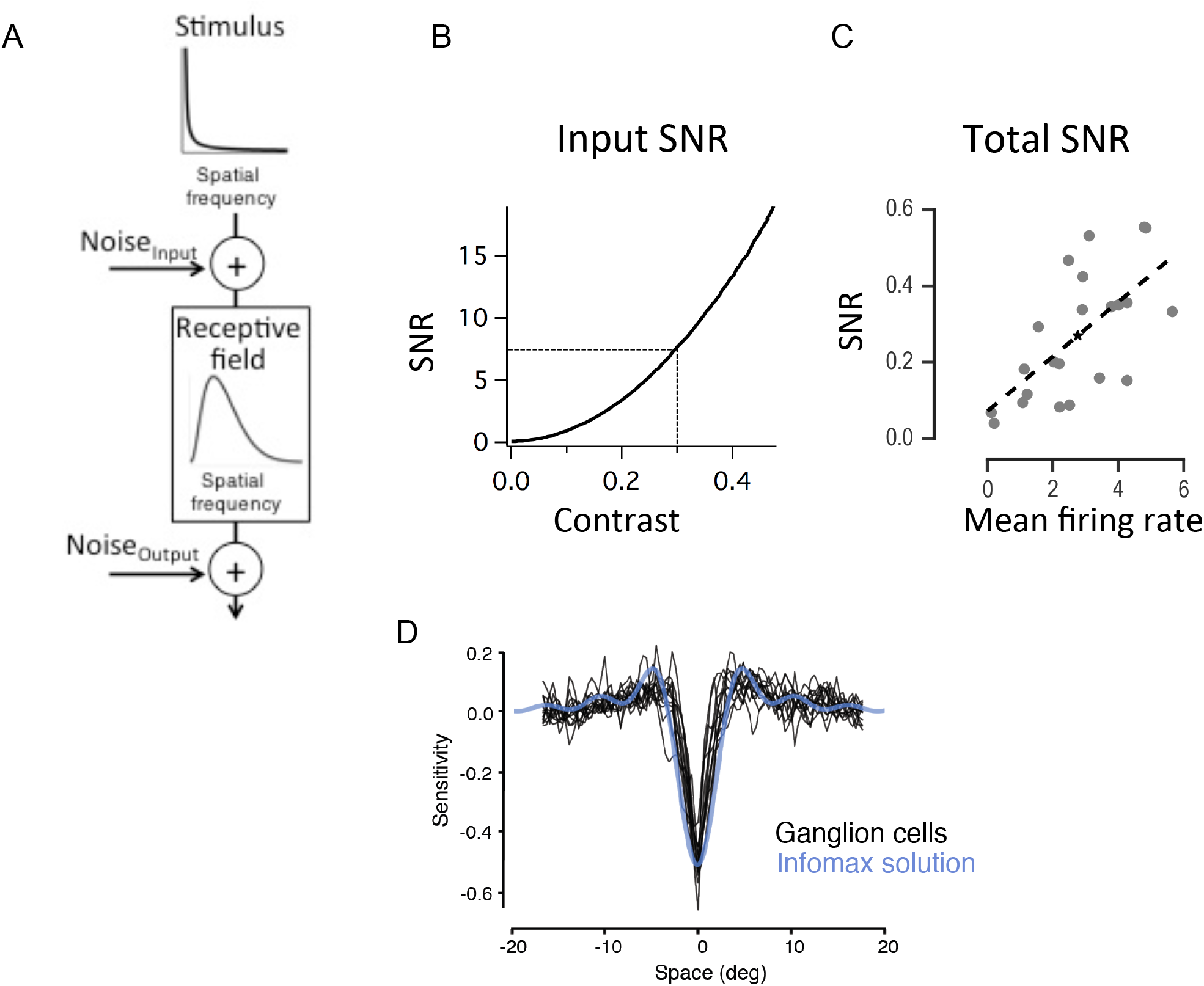
Ganglion cell linear receptive fields maximize information about natural scenes. A) Schematic of model of information transmission through a linear spatial filter in the presence of input and output noise used for Figure 3^3,4^. B) Estimated SNR of cone photoreceptor output as a function of contrast, defined as the standard deviation of the distribution of light intensities divided by the mean intensity. SNR was computed using a mean vesicle rate^42^ of 750 s^−1^, and an assumption of Poisson release. Dashed lines indicate the average value of Michelson contrast of 0.3 computed from natural images^41^, corresponding to a photoreceptor SNR of ~7. C) Total SNR compared to the mean firing rate of each cell for a population of ganglion cells. This value was used to constrain the output noise in the model. D) Comparison of retinal ganglion cell linear receptive fields (black) and the filter that maximizes the mutual information between natural scenes and filter output, subject to a variance constraint (blue). This is the same analysis as in Figure 3A, but in the space domain. The optimal infomax solution is a “brick wall” filter that is zero everywhere the input noise variance exceeds the signal variance.

**Extended Data Figure 3.**
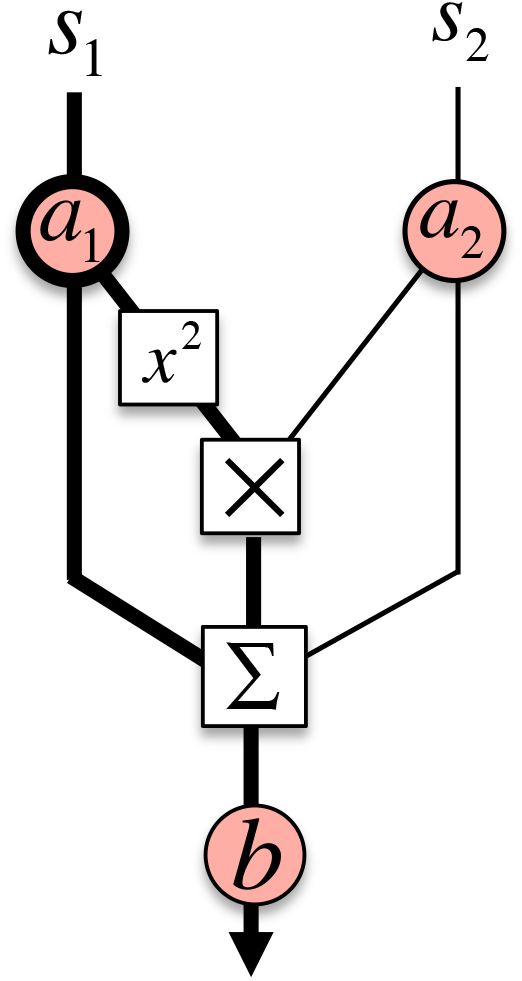
Interneurons can convey a visual feature different than their own receptive field through multiplicative interactions with other pathways. To demonstrate the potential that an interneuron can influence more than its own receptive field component to a downstream neuron, a simple circuit is shown where two stimulus regions *s*_1_ and *s*_2_ are processed by two interneurons *a*_1_ = *s*_1_ and *a*_2_ = *s*_2_ and then contribute to a downstream neuron *b* with an additive and multiplicative interaction, such that 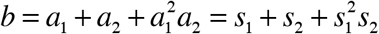. A scaling factor *γ* controls the gain of *a*_1_ as performed experimentally in the record and playback experiment of Fig. 3. Supplementary Information shows that the receptive field component contributed by *a*_1_ includes the receptive fields of both and *a*_2_ by virtue of the multiplicative interaction.

**Extended Data Figure 4.**
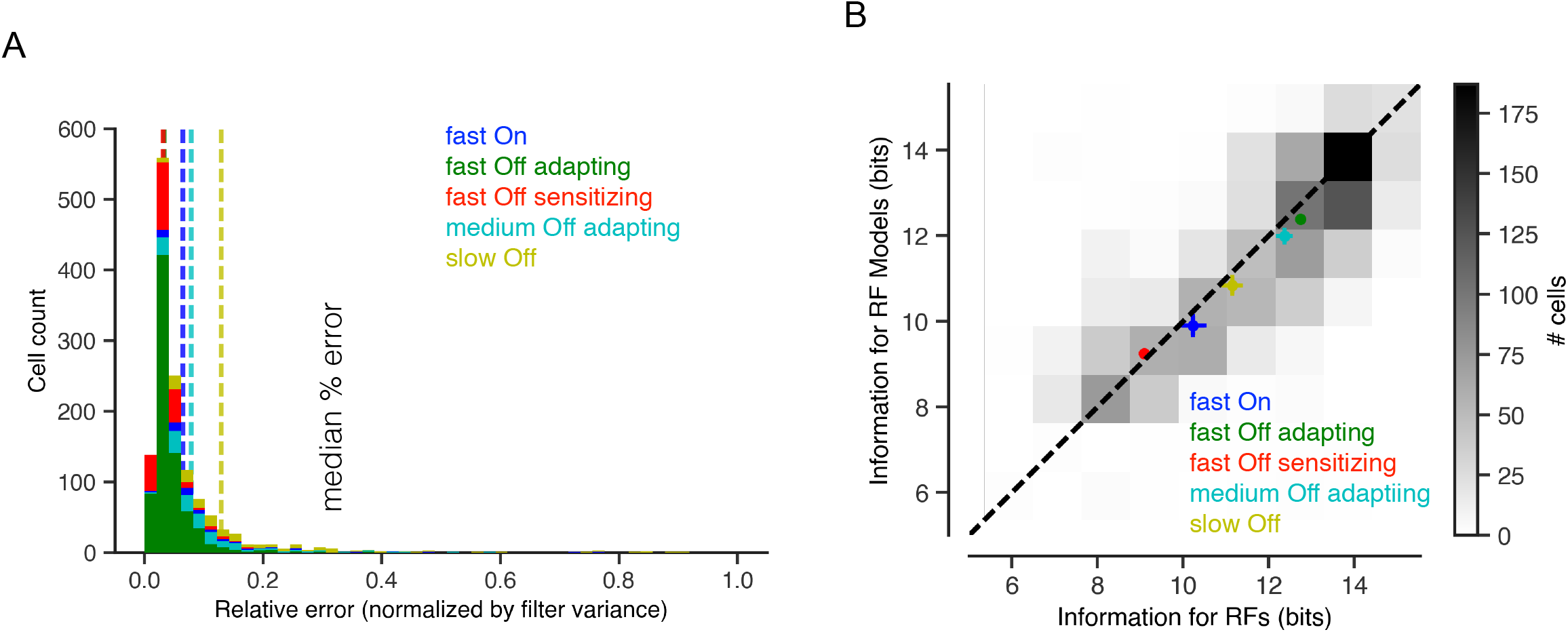
Ganglion cell receptive fields are well fit by linear model of horizontal, amacrine, and excitatory contributions. A) Stacked histogram of the mean squared error between each recorded retinal ganglion cell receptive field and its linear model fit, normalized by the variance of each receptive field. Vertical lines denote the median error for each cell type. B) A two-dimensional histogram of information transmitted by the linear receptive field models versus information transmitted by the receptive fields of the recorded retinal ganglion cells. Points indicate the mean ± s. e. m. for each cell type.

**Extended Data Figure 5.**
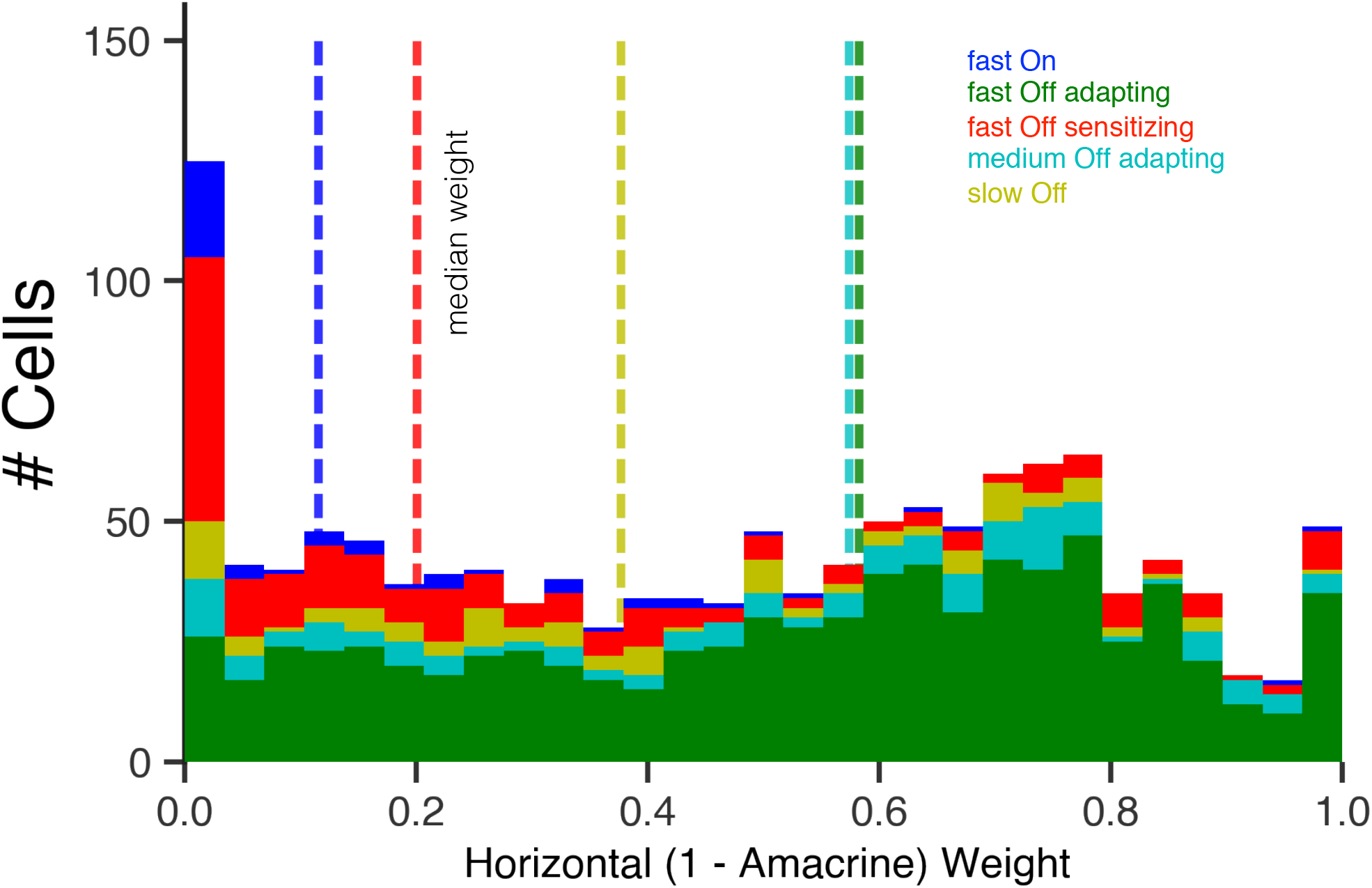
Retinal ganglion cell types have a broad range of horizontal and amacrine cell contributions. Stacked histogram of the relative weighting between horizontal and amacrine cell population contributions to the retinal ganglion cell receptive field, by cell type. Dotted lines denote the median relative horizontal cell weights for each retinal ganglion cell type.

**Extended Data Figure 6.**
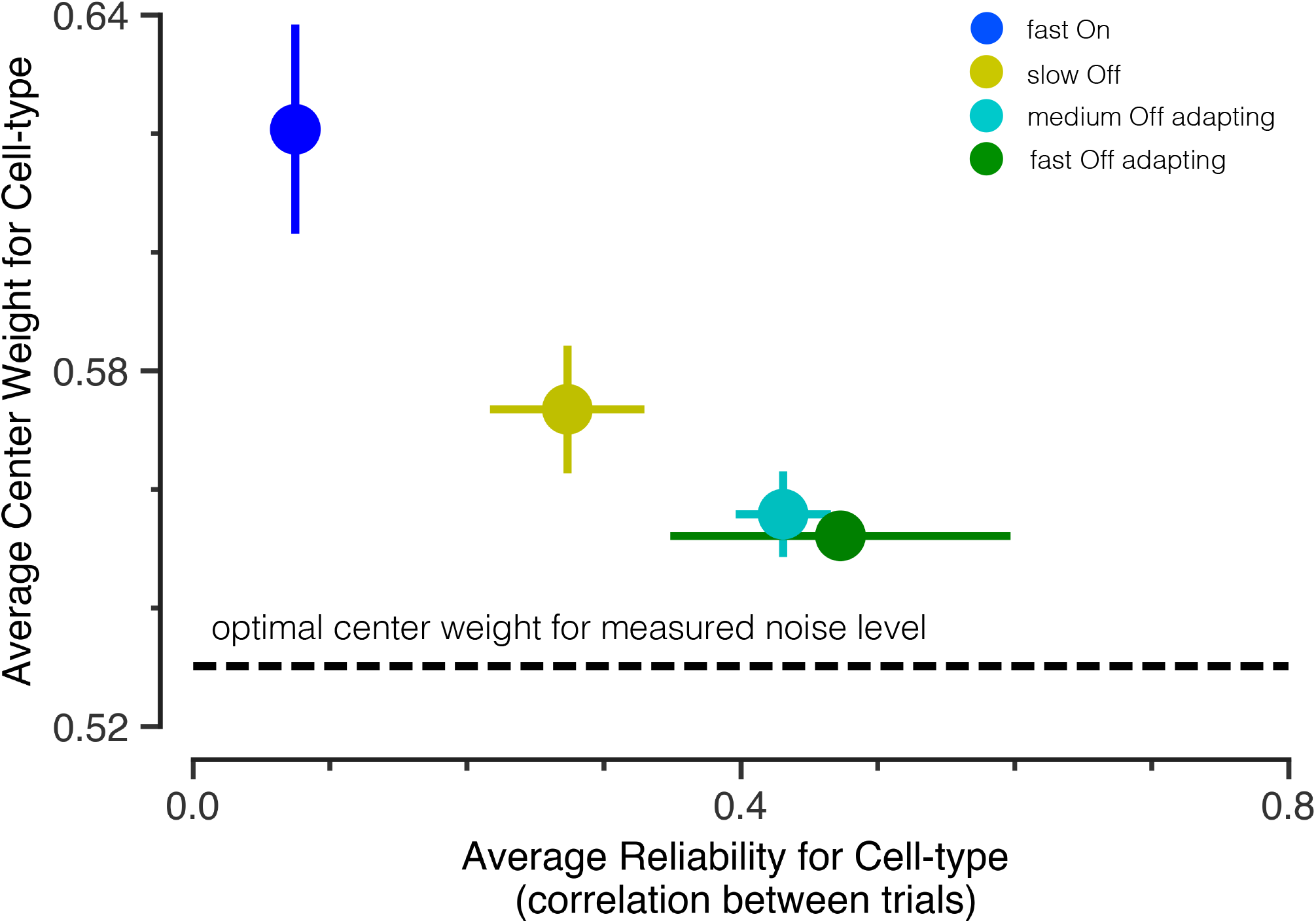
Noisier cell types have weaker surrounds. The average relative center vs surround weight for various retinal ganglion cell types as a function of the trial-to-trial reliability of that cell type measured from a separate dataset of 90 cells responding to a 5 minute 35% contrast white noise repeating stimulus. Cell types that have lower average reliability have weaker surrounds, as measured by an increased center weighting in the linear receptive field model fit. This finding across a population is in accordance with previous work^3^ showing that the optimal receptive field for a lower SNR has a weaker surround.

**Extended Data Figure 7.**
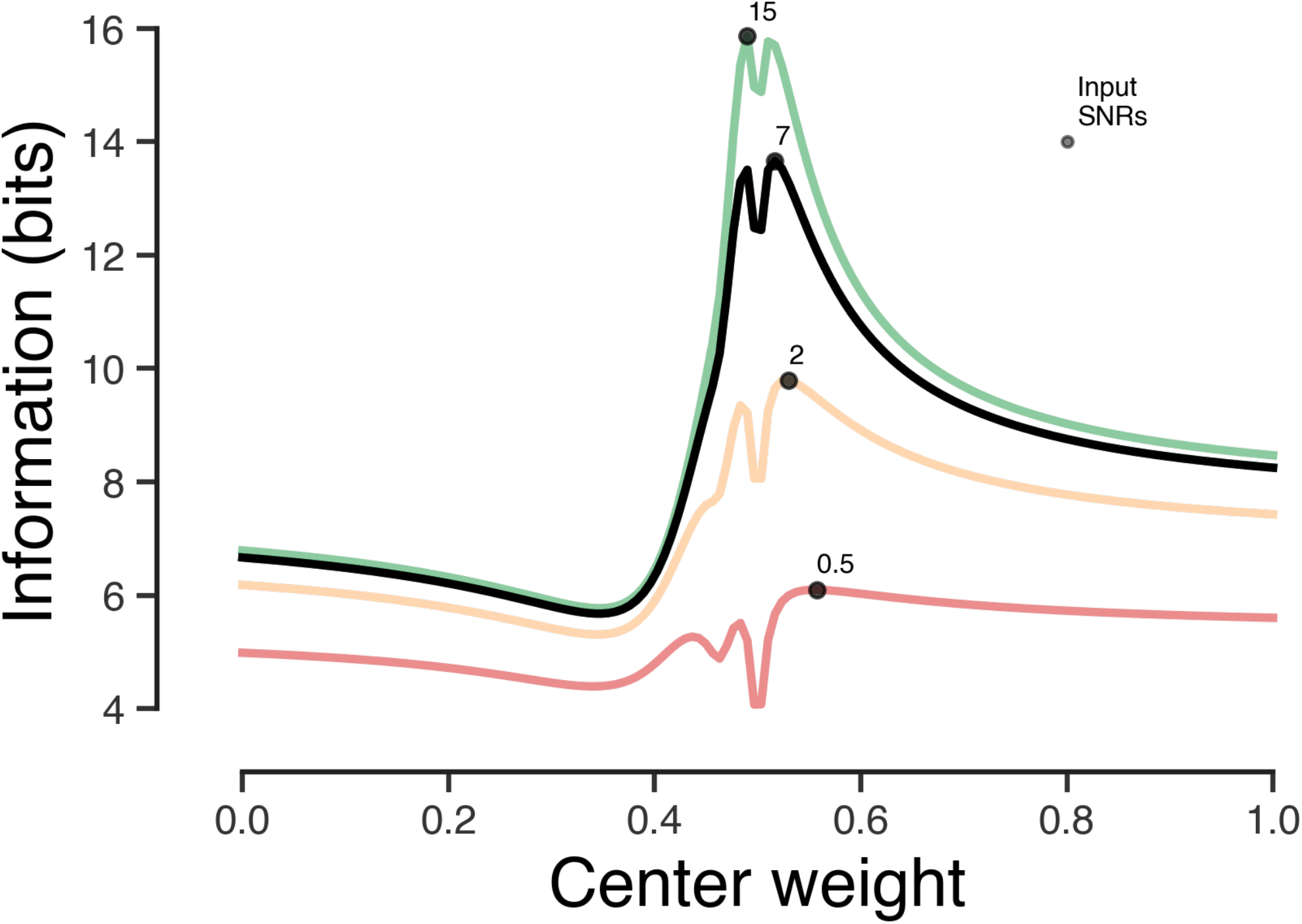
Effect of varying input noise on information landscape. As input noise changes, the ideal information transmitting filter changes shape^3,4^. To quantify how this change in noise effects the qualitative ridge-like appearance of the information landscape in Figure 3D, we varied the input signal-to-noise ratio (colored curves). At low SNRs, the ideal center strength is still qualitatively very similar to what we report in Figure 3, although slightly more monophasic receptive fields are preferred. At an SNR of 15, the optimal center strength moves slightly to the left of the 0.5 center width, indicating a stronger surround relative to the center.

## Contributions

MM, LTM, DBK and SAB designed the study, MM, DBK and BN performed experiments, LTM performed analysis related to optimality of receptive fields, and MM, LTM, DBK, BN and SAB wrote the manuscript.

## Acknowledgments

This work was supported by grants from the NEI, Pew Charitable Trusts, McKnight Endowment Fund for Neuroscience, the Karl Kirchgessner Foundation, and the Alfred P. Sloan Foundation (SAB); by the Stanford Medical Scientist Training Program, NSF IGERT graduate fellowships (LTM, DBK), and an NRSA (BN).

## References

1. Kuffler, S. W. Discharge patterns and functional organization of mammalian retina. J. Neurophysiol. 16, 37–68 (1953).

2. Barlow, H. B. Summation and inhibition in the frog’s retina. J. Physiol. (Lond.) 119, 69–88 (1953).

3. Atick, J. J. & Redlich, A. N. Towards a Theory of Early Visual Processing. Neural Comput 2, 308–320 (1990).

4. Van Hateren, J. Real and optimal neural images in early vision. Nature (1992).

5. Kerr, J. N. D. & Denk, W. Imaging in vivo: watching the brain in action. Nat Rev Neurosci 9, 195–205 (2008).

6. Tye, K. M. & Deisseroth, K. Optogenetic investigation of neural circuits underlying brain disease in animal models. Nat Rev Neurosci 13, 251–266 (2012).

7. Hubel, D. H. & Wiesel, T. N. Receptive fields, binocular interaction and functional architecture in the cat’s visual cortex. J. Physiol. (Lond.) 160, 106–154 (1962).

8. Rodieck, R. W. Quantitative analysis of cat retinal ganglion cell response to visual stimuli. Vision Res. 5, 583–601 (1965).

9. Martinez, L. M. A new angle on the role of feedfoward inputs in the generation of orientation selectivity in primary visual cortex. J. Physiol. (Lond.) 589, 2921–2922 (2011).

10. Naka, K. I. & Nye, P. W. Role of horizontal cells in organization of the catfish retinal receptive field. J. Neurophysiol. 34, 785–801 (1971).

11. Mangel, S. C. Analysis of the horizontal cell contribution to the receptive field surround of ganglion cells in the rabbit retina. J. Physiol. (Lond.) 442, 211–234 (1991).

12. McMahon, M. J., Packer, O. S. & Dacey, D. M. The classical receptive field surround of primate parasol ganglion cells is mediated primarily by a non-GABAergic pathway. J. Neurosci. 24, 3736–3745 (2004).

13. Ichinose, T. & Lukasiewicz, P. D. Inner and outer retinal pathways both contribute to surround inhibition of salamander ganglion cells. J. Physiol. (Lond.) 565, 517–535 (2005).

14. Protti, D. A. et al. Inner retinal inhibition shapes the receptive field of retinal ganglion cells in primate. J. Physiol. (Lond.) 592, 49–65 (2014).

15. Lehky, S. R. & Sejnowski, T. J. Network model of shape-from-shading: neural function arises from both receptive and projective fields. Nature (1988).

16. de Vries, S. E. J., Baccus, S. A. & Meister, M. The projective field of a retinal amacrine cell. J. Neurosci. 31, 8595–8604 (2011).

17. Asari, H. & Meister, M. Frontiers | The projective field of single bipolar cells in the retina. Front Neurosci Conference Abstract: … (2010). doi:10.3389/conf.fnins.2010.03.00145/event_abstract

18. Manu, M. & Baccus, S. A. Disinhibitory gating of retinal output by transmission from an amacrine cell. Proc. Natl. Acad. Sci. U.S.A. 108, 18447–18452 (2011).

19. Masland, R. H. The Neuronal Organization of the Retina. Neuron 76, 266–280 (2012).

20. Doi, E. et al. Efficient Coding of Spatial Information in the Primate Retina. J. Neurosci. 32, 16256–16264 (2012).

21. Hosoya, T., Baccus, S. A. & Meister, M. Dynamic predictive coding by the retina. Nature 436, 71–77 (2005).

22. Hammond, P. Cat retinal ganglion cells: size and shape of receptive field centres. J. Physiol. (Lond.) 242, 99–118 (1974).

23. Devries, S. H. & Baylor, D. A. Mosaic Arrangement of Ganglion Cell Receptive Fields in Rabbit Retina. J. Neurophysiol. 78, 2048–2060 (1997).

24. Kim, I.-J., Zhang, Y., Yamagata, M., Meister, M. & Sanes, J. R. Molecular identification of a retinal cell type that responds to upward motion. Nature 452, 478–482 (2008).

25. Baccus, S. A., Olveczky, B. P., Manu, M. & Meister, M. A retinal circuit that computes object motion. J. Neurosci. 28, 6807–6817 (2008).

26. Olshausen, B. A. & Field, D. J. Emergence of simple-cell receptive field properties by learning a sparse code for natural images. Nature 381, 607–609 (1996).

27. Vinje, W. E. & Gallant, J. L. Sparse Coding and Decorrelation in Primary Visual Cortex During Natural Vision. Science 287, 1273–1276 (2000).

28. Padmanabhan, K. & Urban, N. N. Intrinsic biophysical diversity decorrelates neuronal firing while increasing information content. Nat. Neurosci. (2010).

29. Mejias, J. F. & Longtin, A. Optimal Heterogeneity for Coding in Spiking Neural Networks. Phys. Rev. Lett. 108, 228102 (2012).

30. Pitkow, X., Sompolinsky, H. & Meister, M. A Neural Computation for Visual Acuity in the Presence of Eye Movements. PLoS Biol. 5, e331 (2007).

31. Gjorgjieva, J., Sompolinsky, H. & Meister, M. Benefits of Pathway Splitting in Sensory Coding. J. Neurosci. 34, 12127–12144 (2014).

32. Kastner, D. B., Baccus, S. A. & Sharpee, T. O. Critical and maximally informative encoding between neural populations in the retina. Proc. Natl. Acad. Sci. U.S.A. 112, 2533–2538 (2015).

33. Liu, Y. S., Stevens, C. F. & Sharpee, T. O. Predictable irregularities in retinal receptive fields. Proc. Natl. Acad. Sci. U.S.A. 106, 16499–16504 (2009).

34. Baden, T. et al. The functional diversity of retinal ganglion cells in the mouse. Nature 529, 345–350 (2016).

35. van Hateren, J. H. A theory of maximizing sensory information. Biol Cybern (1992).

36. Linsker, R. Local Synaptic Learning Rules Suffice to Maximize Mutual Information in a Linear Network. http://dx.doi.org/10.1162/neco.1992.4.5.691 4, 691–702 (2008).

37. Karklin, Y. & Simoncelli, E. P. Efficient coding of natural images with a population of noisy linear-nonlinear neurons. Advances in neural information processing … (2011).

38. Otchy, T. M. et al. Acute off-target effects of neural circuit manipulations. Nature 528, 358–363 (2015).

39. Kastner, D. B. & Baccus, S. A. Coordinated dynamic encoding in the retina using opposing forms of plasticity. Nat. Neurosci. 14, 1317–1322 (2011).

40. Chichilnisky, E. J. A simple white noise analysis of neuronal light responses. Network 12, 199–213 (2001).

41. Tkačik, G. et al. Natural Images from the Birthplace of the Human Eye. PLoS ONE 6, e20409 (2011).

42. Devries, S. H., Li, W. & Saszik, S. Parallel Processing in Two Transmitter Microenvironments at the Cone Photoreceptor Synapse. Neuron 50, 735–748 (2006).

43. Tadmor, Y. & Tolhurst, D. J. Calculating the contrasts that retinal ganglion cells and LGN neurones encounter in natural scenes. Vision Res. 40, 3145–3157 (2000).

